# A cytoplasmic protein kinase in *Chlamydomonas* couples engagement of ciliary receptors to rapid cellular responses

**DOI:** 10.1101/2021.09.03.458889

**Authors:** Mayanka Awasthi, Peeyush Ranjan, Simon Kelterborn, Peter Hegemann, William J. Snell

## Abstract

The principal function of the primary cilium is to convert cues from the extracellular milieu into changes in cyclic nucleotide concentration and cytoplasmic responses, but fundamental questions remain about the mechanisms of transmission of cilium-to-cytoplasm signals. During fertilization in *Chlamydomonas reinhardtii*, ciliary adhesion between *plus* and *minus* gametes triggers an immediate ∼10-fold increase in cellular cAMP and activation for cell fusion. Here, we identify <u>G</u>amete-<u>S</u>pecific <u>P</u>rotein <u>K</u>inase (GSPK) as an essential link between cilary receptor engagement and gamete activation. The ciiary adhesion-induced increase in cAMP and cell fusion are severely impaired in *gspk* mutants but fusion is rescued by a cell-permeable form of cAMP, indicating that GSPK functions upstream of the cAMP increase. GSPK is cytoplasmic, and, remarkably, the entire cellular complement is phosphorylated in less than 60 seconds after ciliary contact. Thus, a cytoplasmic protein kinase rapidly converts a ciliary membrane cue into a global cellular response.

## Introduction

The primary cilium is a spatially distinct cellular compartment specialized for receipt of extracellular signals. Its membrane houses multiple receptors essential for development and homeostasis^1-3^, including G-protein coupled receptors (GPCRs) responsible for vision^4^, olfaction^5,6^, growth factor receptor-activated pathways^7^, regulation of insulin and glucagon secretion in pancreatic islet cells^8^, and the sonic hedgehog (Hh) developmental pathway^9^. And, cilia participate in regulation of epithelial cell proliferation in the liver, gall bladder, and kidney^3,10,11^. The ciliary compartment is unique in its combined functional and physical separation from the cell proper and is surrounded, not by cytoplasm, as are intracellular membrane-bounded organelles, but by the extracellular milieu. Moreover, the membrane at the connection between the cilium and the cell is packed with membrane protein complexes tightly linked to the dense array of underlying doublet microtubules of the ciliary axoneme^12-14^, thereby providing a regulatable barrier to membrane protein movement between the two domains.

Although the multiple signaling pathways initiated in the cilium vary widely across cell types in their mechanisms of activation and in their downstream outcomes, almost all are linked by their common function in regulating the concentrations of cyclic nucleotides, primarily the second messenger, cAMP. Increases in ciliary cAMP upon odorant binding by cilia of olfactory epithelial cells alter the activity of cyclic nucleotide-gated ion channels within the cilia, thereby inducing changes in membrane potential that are transmitted through the cell body and axons to generate a sense of smell^15^. Changes in ciliary cAMP upon activation of the cilium-based sonic hedgehog (Hh) and other GPCR-regulated pathways alter the activities of cAMP-dependent protein kinases and EPACs through multiple, complex mechanisms that lead to changes in protein secretion and gene expression^16-18^. During kidney tubule development, the increased cAMP that is a consequence of mutation of ciliary proteins polycystin I, polycystin 2, fibrocystin, and others leads to polycystic kidney disease (PKD)^3,11^. In spite of the importance of cilium-initiated, cAMP-dependent signaling pathways, however, fundamental questions remain about the mechanisms that couple receptor activity at the ciliary membrane to downstream responses in the cell.

In many unicellular organisms, including parasitic protozoans, ciliated protozoans, and green algae, cues received at cilia trigger changes in ciliary cAMP and cellular responses, and thus this ciliary signaling strategy is ancient^19-23^. During sexual reproduction in the bi-ciliated, unicellular green alga *Chlamydomonas reinhardtii*, interactions between the cilia of gametes of opposite mating types trigger a rapid, ∼10-fold increase in intracellular cAMP that activates the gametes for cell-cell fusion^22-26^. Ciliary adhesion is mediated by binding between adhesion receptor SAG1 on the cilia of *plus* gametes and adhesion receptor SAD1 on the cilia of *minus* gametes^27,28^. During this flirtation with multicellularity by a unicellular organism, the signaling pathway activated by SAG1-SAD1 interactions exhibits many of the hallmarks of pathways activated by receptor-ligand interactions in animal cells, including changes in the phosphorylation states of proteins and changes in protein location^23-26,29,30^. *Chlamydomonas* ciliary signaling has proved useful for understanding fundamental signaling properties of cilia. Studies over 35 years ago demonstrated the existence of a functional barrier between the *Chlamydomonas* plasma membrane and the ciliary membrane^31,32^. Related studies showed that, contrary to then emerging models, regulated movement of membrane proteins into the cilia does not require intraflagellar transport (IFT)^33-35^.

Studies with cilia isolated separately from naive (resting or unmixed) *plus* and *minus* gametes and from gametes soon after mixing have shown that within seconds after cilia adhere to each other, a cGMP-dependent protein kinase (PKG) becomes phosphorylated^36,37^. Although the protein kinase that phosphorylates the PKG and the substrates for the PKG are unknown, experimentally reducing expression of the PKG impairs gamete fusion^37^. Related studies have also shown that cilia isolated from *Chlamydomonas* gametes possess an adenylyl cyclase activity that is regulated by phosphorylation and dephosphorylation^25,26^. The in vitro activity of this as yet unidentified ciliary adenylyl cyclase is increased ∼2-fold after *plus* and *minus* gametes are mixed together^24^, and even when isolated cilia are mixed together^26^. Moreover, cell bodies possess an adenylyl cyclase activity detected by in vitro assays, that is also increased ∼2-fold by ciliary adhesion. The adhesion-induced increase in the ciliary adenylyl cyclase activity occurs within 1 minute after gametes are mixed together, and the increase in the activity of the cell body adenylyl cyclase occurs within ∼2 minutes. Although, one model is that the cAMP formed in the cilia activates the adenylyl cyclase in the cell body^24^, the relationship between the ciliary adenylyl cyclase activity and gamete activation is unknown.

The primary cellular responses to the ciliary adhesion-triggered increase in cellular cAMP occur in the cell body within ∼2 minutes after gametes are mixed together. The gametes release their cell walls^38^, mobilize pools of SAG1 and SAD1 from the plasma membrane onto the ciliary membrane in a positive feedback mechanism that sustains and enhances adhesion,^30,34,39-41^ and erect fusogenic membrane protuberances - - mating structures^42^- - as the gametes prepare for fusion to form a quadri-ciliated zygote (Fig. 1a). These cellular responses to ciliary adhesion (with the exception of gamete fusion) can be mimicked experimentally by incubation of gametes of a single mating type in a buffer containing a cell-permeable analogue of cAMP, dibutyryl cAMP (db-cAMP)^22^. As in other systems, however, the molecules and mechanisms that couple ciliary receptor engagement in *Chlamydomonas* gametes to cAMP-dependent cellular responses in the cytoplasm remain poorly understood.

**Figure 1.**
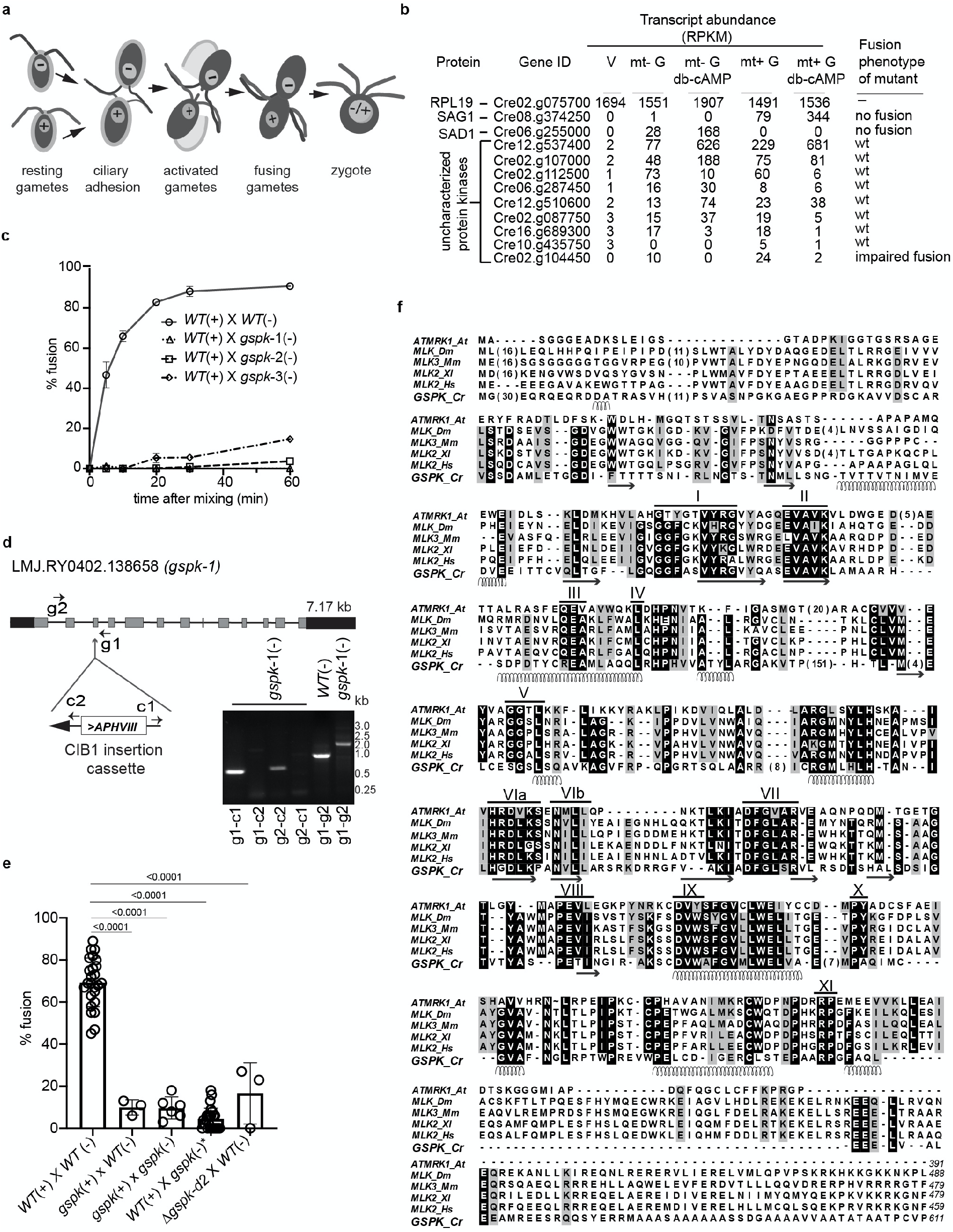
GSPK is a gamete-specific mixed lineage protein kinase essential for gamete fusion. **a** Illustration of fertilization in *Chlamydomonas*. **b** The Cre02.g104450 mutant has a fusion phenotype. Transcript abundance for the indicated genes in vegetatative (V), naive *minus* (mt-G) and *plus* (mt+ G) gametes, and db-cAMP-activated *minus* (mt-G, db-cAMP) and *plus* (mt+ G, db-cAMP) gametes represented by median reads per kilobase per million mapped reads (RPKM) are from Ning et al. (2013). *RPL19* is a housekeeping gene. The right panel indicates the fusion phenotypes. **c** Fusion is impaired in multiple Cre02.g104450 mutant strains. Quantification of fusion of *WT*(+) gametes with *minus* gametes of *WT* and three different CLiP *gspk* mutants. Values are means (+/- standard deviation) from three replicates. **d** Structure of the *GSPK* genomic locus in the *gspk-1* CLiP mutant. Grey solid boxes indicate exons; thin lines, introns; UTRs are shown as black boxes. The ClB1 (*AphVIII)* insertion cassette is in the 3rd intron of *gspk* in the *gspk* allele of LMJ.RY0402.138658_1 strain. Primer locations for genotyping are shown by arrows, where g1 and g2 are gene-specific primers and c1 and c2 are specific to the insertion cassette. Gel images show diagnostic genomic DNA PCR samples. Primer combinations used for PCR are indicated at the bottom of the lanes. **e** Quantification of fusion in *WT* gametes (n=25 experiments) compared with fusion in *gspk*(*+*) mutant progeny generated from a cross between *gspk*(-) and *WT*(+) (n=3); fusion of *gspk*(*+*) mutants mixed with *gspk*(-) (n=7); fusion of *gspk\**(-) mutant progeny generated from a cross between *gspk*(+) and *WT*(-) (n=25); and fusion of CRISPR-generated Δ*gspk-d2*(+) mutant mixed with *WT*(-) (n=3). P-values labelled above the groups compared are from Student’s t-test. (F) Multiple sequence alignment of GSPK and protein kinases from *Arabidopsis thaliana* (BAA22079.1), *Drosophila melanogaster* (AAL08011.1), *Mus musculus* (AAF73281.1), *Xenopus leavis* (AAP46399.1), and *Homo sapiens* (CAA62351.1). Identical and similar residues are highlighted in black and grey, respectively. The conserved domains of the catalytic core are indicated by roman numerals. Regions with α-helices (spirals) and β-sheets (arrows) predicted for GSPK by JPred4 are denoted below the sequences.

Here, we report identification of a <u>g</u>amete-specific protein kinase, <u>G</u>amete <u>S</u>pecific <u>P</u>rotein <u>K</u>inase (GSPK), that is essential for this cilium-based signaling pathway. GSPK has sequence homology to mixed lineage protein kinases of animal cells and cell fractionation shows that it is a cell body protein. GSPK is basally phosphorylated in naive gametes and undergoes further phosphorylation within 1 minute after *plus* and *minus* gametes are mixed together. Studies with *gspk* mutants indicate that the protein is essential for gamete fusion and functions downstream of ciliary adhesion and the adhesion-induced phosphorylation of ciliary PKG. Cell body responses to ciliary adhesion are strongly impaired in *gspk* gametes, and the rapid cell-cell fusion that typifies wild type gametes fails to occur in the mutants. Experiments showing rescue of fusion in mutant gametes by db-cAMP indicate that the downstream gamete activation machinery is intact in the mutants. Importantly, assays of cellular cAMP indicate that ciliary adhesion in the mutant gametes fails to induce the large increase in cAMP that typically accompanies ciliary adhesion. Our results indicate that GSPK is a cytoplasmic protein that rapidly detects ciliary adhesion and couples engagement of ciliary adhesion receptors to cAMP-dependent responses in the cytoplasm required for cell-cell fusion.

## Results

### Identification of a gamete-specific protein kinase essential for fertilization in *Chlamydomonas*

To identify protein kinases with a potential role in ciliary signaling during fertilization, we tested for a cell-cell fusion phenotype in several *minus* mating-type strains from the *Chlamydomonas* CLiP mutant library that were annotated to contain mutations in protein kinase genes and that exhibited gamete-specific expression profiles^43^ (Fig. 1b). Of 9 strains examined, gametes prepared from strain LMJ.RY0402.138658, which was annotated to have an insertion of the antibiotic resistance cassette *APHVIII* in gene Cre02.g104450 (whose encoded protein is GSPK), was strongly impaired in gamete fusion when mixed with wild type (*WT*) *plus* gametes (Fig. 1b, c). PCR analyses using gene-specific and cassette-specific sets of primers confirmed *APHVIII* cassette insertion into exon 3 in strain LMJ.RY0402.138658 (*gspk-1*) (Fig. 1d and Supplementary Table 1) and at the predicted insertion sites in two other independent CLiP library strains annotated to have insertions in Cre02.g104450, LMJ.RY0402.097798 (*gspk*-2; predicted insertion in exon 8) and LMJ.RY0402.039382 (*gspk*-3; predicted insertion in the 3’UTR) (Supplementary Table 1 and Supplementary Fig. 1). Determination of the percent of gametes that fused after mixing with *WT plus* gametes confirmed that GSPK indeed was essential for the rapid fusion that typifies *WT* gametes. Nearly 70% of the cells in mixtures of *WT minus* gametes and *WT plus* gametes had fused to form quadri-ciliated cells (zygotes) within 10 minutes after mixing (Fig. 1c), whereas, fusion was less than 1% for each of the three *minus* mutant strains (Fig. 1c). The percent fusion in the mutants increased slightly by 60 minutes‘(to 15-18%) (Fig. 1c).

Analysis of progeny from zygotes produced by crossing (Supplementary Fig. 2) *gspk-1 minus* gametes with *WT plus* gametes showed that both *minus* gametes and *plus* gametes bearing the *gspk-1* allele were defective in fusion when mixed with *WT* gametes of the opposite mating type (Fig. 1e). The identical phenotype was also found in a separate Cre02.g104450 mutant strain Δ*gspk-d2* generated in *plus* cells by use of CRISPR methods (Fig. 1e; Supplementary Fig. 3). Thus, the *gspk* mutation segregated with the mutant phenotype, and *gspk* gametes of both mating types exhibited the fusion phenotype.

Examination of sequence alignments showed that GSPK possessed the canonical protein kinase sub-domains of members of the protein kinase superfamily and was most closely related to mixed lineage protein kinases (Fig. 1f). GSPK contains ∼151 residues between subdomains IV and V absent in most other protein kinases that could be a potential regulatory motif. Use of NMT - The MYR Predictor (https://mendel.imp.ac.at/myristate/SUPLpredictor.htm) to predict N-myristoylation sites indicated that the glycine at position 2 in GSPK (M<u>G</u>AVLSCCGEGTIGASHG) is a potential myristoylation site.

To investigate the cellular properties of GSPK, we introduced into *gspk* cells a transgene encoding an epitope-tagged form of GSPK, *GSPK-HA*, driven by the endogenous promoter. Immunoblotting of *gspk* and *gspk*/*GSPK-HA minus* gametes with anti-HA antbodies showed a tagged protein of the expected size, ∼70 kDa, only in the cells bearing the transgene (Fig. 2a). Consistent with the analysis above indicating that mutation of *GSPK* was responsible for the fusion phenotype, introduction of the *GSPK-HA* transgene rescued fusion (Fig. 2a). Furthermore, and consistent with the transcriptome evidence (Fig. 1b), GSPK-HA was expressed only in gametes and not vegetative cells, and activation of the gametes by incubation in db-cAMP buffer for 1 hour brought about a substantial reduction in GSPK-HA protein levels (Fig. 2b).

**Figure 2:**
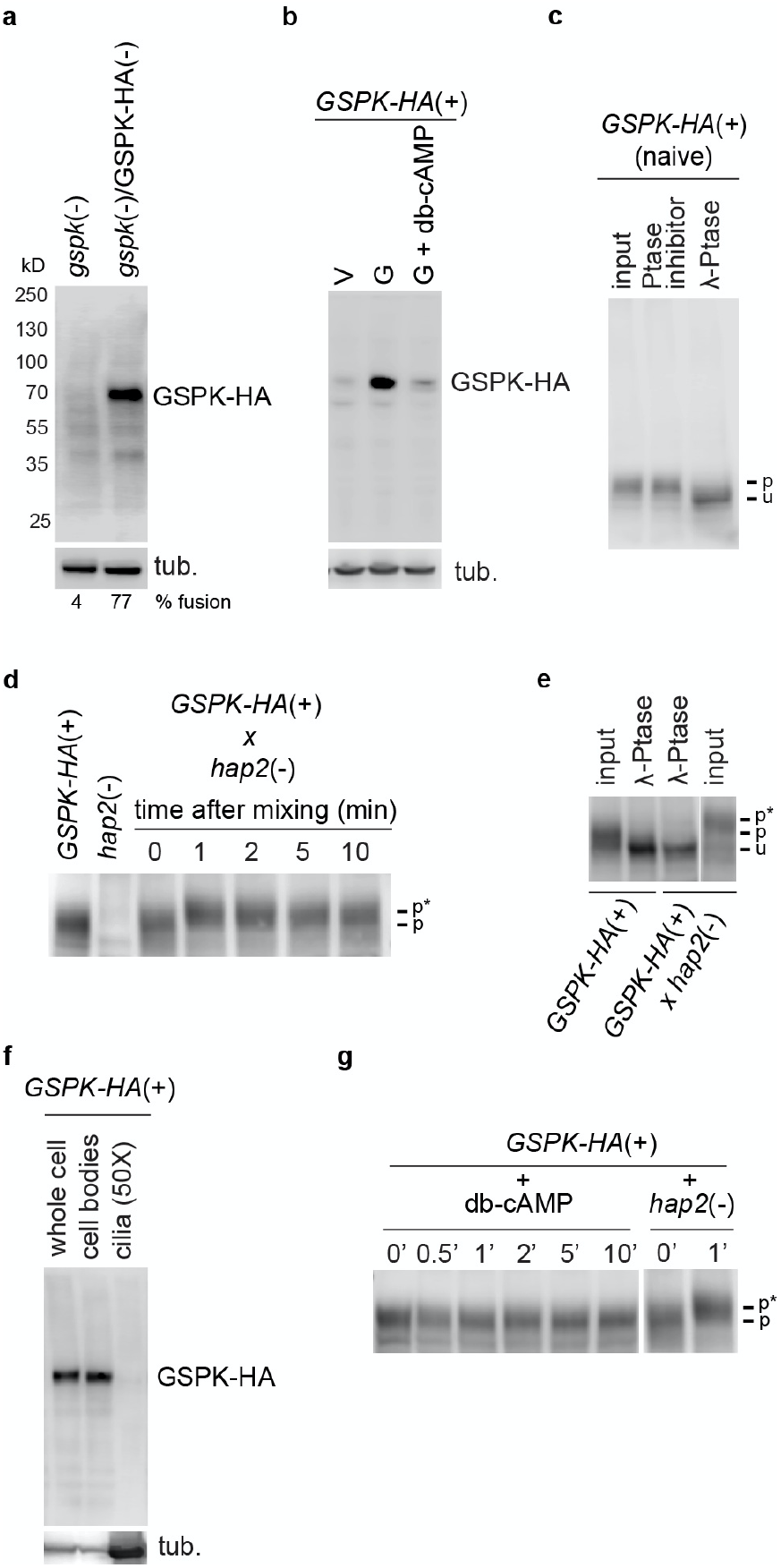
GSPK is gamete-specific and phosphorylated within 1 minute after ciliary adhesion is initiated. **a** Expression of GSPK-HA in *gspk minus* gametes rescues fusion. Anti-HA immunoblot of *gspk*(-) and *GSPK-HA*-rescued *gspk*(-) gametes. Percent fusion at 10 minutes is shown below the blot. Tubulin was used as a loading control. **b** GSPK-HA expression is gamete-specific and reduced upon gamete activation by db-cAMP. Anti-HA immunoblot of *gspk/GSPK-HA*(+) vegetative cells (V), naive gametes (G), and gametes activated by incubation in db-cAMP buffer (G-A). **c** GSPK-HA is basally phosphorylated in naive gametes. Anti-HA immunoblot of lysates of *gspk/GSPK-HA*(+) naive gametes that had been incubated with λ-phosphatase in the presence and absence of a phosphatase inhibitor. **d-e** GSPK-HA is phosphorylated within 1 minute after initiation of ciliary adhesion. Anti-HA immunoblots of *GSPK-HA*(+) gametes at the indicated times after mixing with *hap2* (-) gametes (**d**). Anti-HA immunoblot of lysates of *gspk/GSPK-HA*(+) gametes before and 10 minutes after mixing with *hap2*(-) gametes with and without treatment with λ-phosphatase (**e**). Letters on the right indicate GSPK-HA that is unphosphorylated (u), basally phosphorylated (p), or additionally phosphorylated (p*). **f** GSPK fractionates with cell bodies. Anti-HA immunoblot of whole cells, cell bodies, and cilia of *gspk/GSPK-HA*(+) gametes. 3 μg of protein were loaded per lane, which for cilia represents about 50 cell equivalents. The lower panel is a tubulin loading control. **g** Activation of gametes with db-cAMP buffer fails to induce phosphorylation of GSPK-HA. Anti-HA immunoblot of *gspk/GSPK-HA*(+) gametes at the indicated times after mixing with db-cAMP buffer. Cell wall loss was over 80% at 10 minutes, confirming gamete activation.

### The entire cellular complement of GSPK is in the cytoplasm and is phosphorylated within 1 minute after ciliary receptor engagement

We used SDS-PAGE and immunoblotting to assess the phosphorylation state of GSPK-HA in naive *plus* gametes and in *plus* gametes undergoing ciliary adhesion at increasing times after mixing with *hap2 minus* gametes. As shown in Fig. 2c, incubation of lysates of naive *GSPK-HA* gametes with the protein de-phosphorylating enzyme, l-phosphatase, led to a shift in migration of GSPK-HA compared to the non-treated sample or compared to a sample incubated with the phosphatase and a phosphatase inhibitor. These results indicated that GSPK was basally phosphorylated in naive gametes.

Similar analysis showed that ciliary adhesion induced a further increase in GSPK-HA phosphorylation. Upon mixing the *GSPK-HA*(+) gametes with *hap2 minus* gametes, which are defective in gamete fusion because they fail to express the gamete fusogen, HAP2^44^, the basally phosphorylated GSPK-HA underwent a shift in migration (Fig. 2d). Remarkably, the entire cellular complement of GSPK-HA underwent the shift, and the shift was detectable within 1 minute after mixing. Consistent with the shift being a consequence of phosphorylation, all of the GSPK-HA was shifted to the unphosphorylated form upon incubation of the lysates with l-phosphatase (Fig. 2e).

Given that all of the GSPK underwent the ciliary adhesion-induced rapid phosphorylation, it seemed likely that the protein itself would be localized in the organelles. Analysis by immunoblotting, however, of naive whole cells, cell bodies, and cilia indicated that GSPK was present in cell bodies, with little if any detectable in the cilia (Fig. 2f). Moreover, even though all other cell body events that occur during gamete interactions can be induced in gamete of a single mating type by incubation in db-cAMP^32,41^, phosphorylation of GSPK-HA was not induced by db-cAMP (Fig. 2g). (Cell wall loss was over 80% at 10 minutes in these samples). Thus, interactions between SAG1 and SAD1 at the surface of the cilia were rapidly transduced into phosphorylation of GSPK in the cell body, but GSPK phosphorylation was upstream of the increase in cAMP that drives gamete activation.

### The earliest biochemically detectable response in cilia to ciliary adhesion, phosphorylation of a cGMP-dependent protein kinase, does not require GSPK

To examine the cellular function of GSPK, we further investigated the phenotype of *gspk* mutants. Vegetative cells of both *plus* and *minus gspk* strains were indistinguishable from *wild-type* vegetative cells in size, appearance, motility, and growth. Moreover, all of the *gspk* mutant strains underwent normal gametogenesis to form gametes that were indistinguishable from the wild-type gametes in morphology and motility (not shown). Similarly, microscopic examination (Fig. 3ai) and a quantitative assay for ciliary adhesion (Fig. 3aii) showed that the *gspk minus* gametes underwent initial ciliary adhesion with *WT plus* gametes to nearly the same extent as did fusion-defective *hap2 minus* gametes with *WT plus* gametes. *gspk plus* gametes were similarly competent for ciiary adhesion with *WT minus* gametes (not shown). Thus, GSPK functioned downstream of SAG1-SAD1-dependent ciliary adhesion.

**Figure 3:**
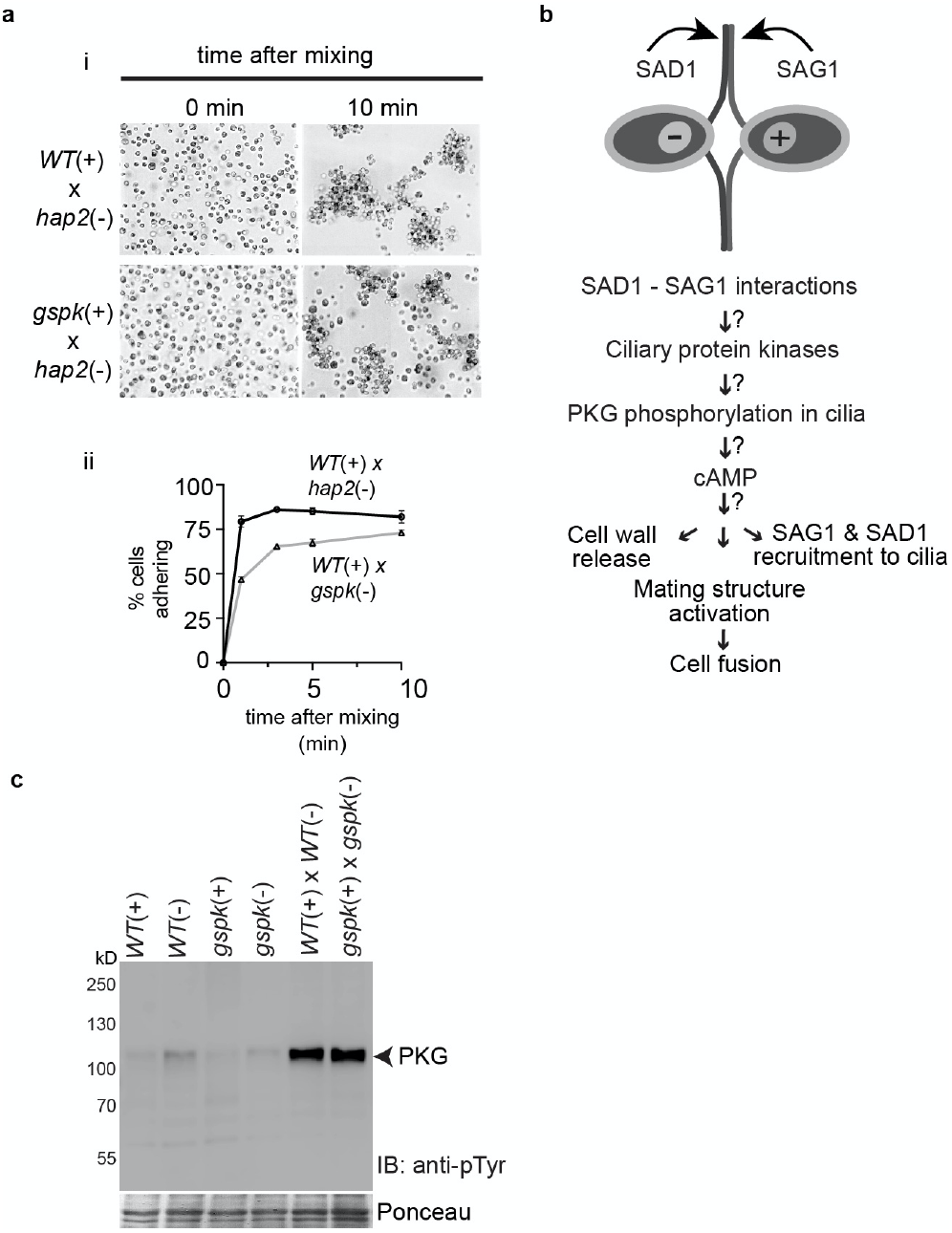
*gspk* mutants undergo ciliary adhesion and phosphorylation of PKG in cilia similarly to *WT*. **a** *gspk*(+) gametes undergo normal ciliary adhesion. **ai** Differential Interference Contrast (DIC) micrographs of *WT*(+) gametes (upper panel) and *gspk*(+) gametes (lower panel) at the times indicated after mixing with *hap2*(-) gametes. **aii** Quantification of adhesion in the indicated samples by use of an electronic particle counter. **b** Schematic diagram of cilium-generated signaling pathway in *Chlamydomonas*. **c** Ciliary adhesion in *gspk* gametes induces phosphorylation of cGMP-dependent protein kinase (PKG). Anti-p-Tyr immunoblots from PTK assay of cilia isolated from non-adhering *WT*(+), *WT*(-), *gspk*(+) and *gspk*(-) gametes and from *WT*(+) gametes mixed with *WT*(-) gametes and *gspk*(+) gametes mixed with *gspk*(-) gametes.

We tested whether the earliest experimentally detectable consequence of ciliary adhesion, phosphorylation of ciliary PKG (Fig. 3b), was intact in the *gspk* mutants. Cilia isolated from *WT plus* and *minus* gametes that had been mixed together for 3 minutes and from *gspk* mutant *plus* and *minus* gametes mixed for the same time were assessed for tyrosine phosphorylation of PKG by an in vitro assay and immunoblotting with anti-phosphotyrosine antibodies^36^. As shown in Fig. 3c, phosphorylation of the 105 kDa PKG was at very low levels in assays of cilia isolated from naive *plus* gametes and from naive *minus WT* gametes and in assays of separately isolated cilia from naive *plus* and *minus gspk* gametes. On the other hand, cilia isolated from the mixed *WT plus* and *minus* gametes and from the mixed *gspk plus* and *minus* gametes undergoing ciliary adhesion showed robust phosphorylation of PKG in the assays. Thus, the earliest biochemical response within cilia to SAG1-SAD1 interactions was independent of GSPK.

### Ciliary adhesion by *gspk* gametes fails to induce the cell body responses required for gamete fusion

Given that the block to fusion in the *gspk* mutants was downstream of initial ciliary events, we used bioassays to determine whether *gspk* gametes underwent the typical cell body responses to ciliary adhesion (Fig. 3b). Our wall loss assay indicated that cell wall release was severely impaired in the *gspk* gametes (Fig. 4a). Whereas nearly 70% of the cells in samples of adhering *WT plus* gametes mixed with *hap2 minus* gametes had lost their walls at 10 minutes after mixing, fewer than 20% of the mixed *gspk* gametes had lost their walls. Similary, mating structure activation, as measured by the appearance of the actin-filled microvillous-like fertilization tubules in *plus* gametes was substantially reduced in the *gspk* gametes (Fig. 4b). 30 minutes after mixing equal numbers of *WT plus* gametes with *hap2 minus* gametes, nearly 45% of the cells in the mixture possessed actin-staining mating structures (which meant that ∼90% of the *plus* gametes had formed mating structure), but fewer than 3% of the *gspk plus* gametes had formed the structures (Fig. 4b and Supplementary Fig. 4).

**Figure 4:**
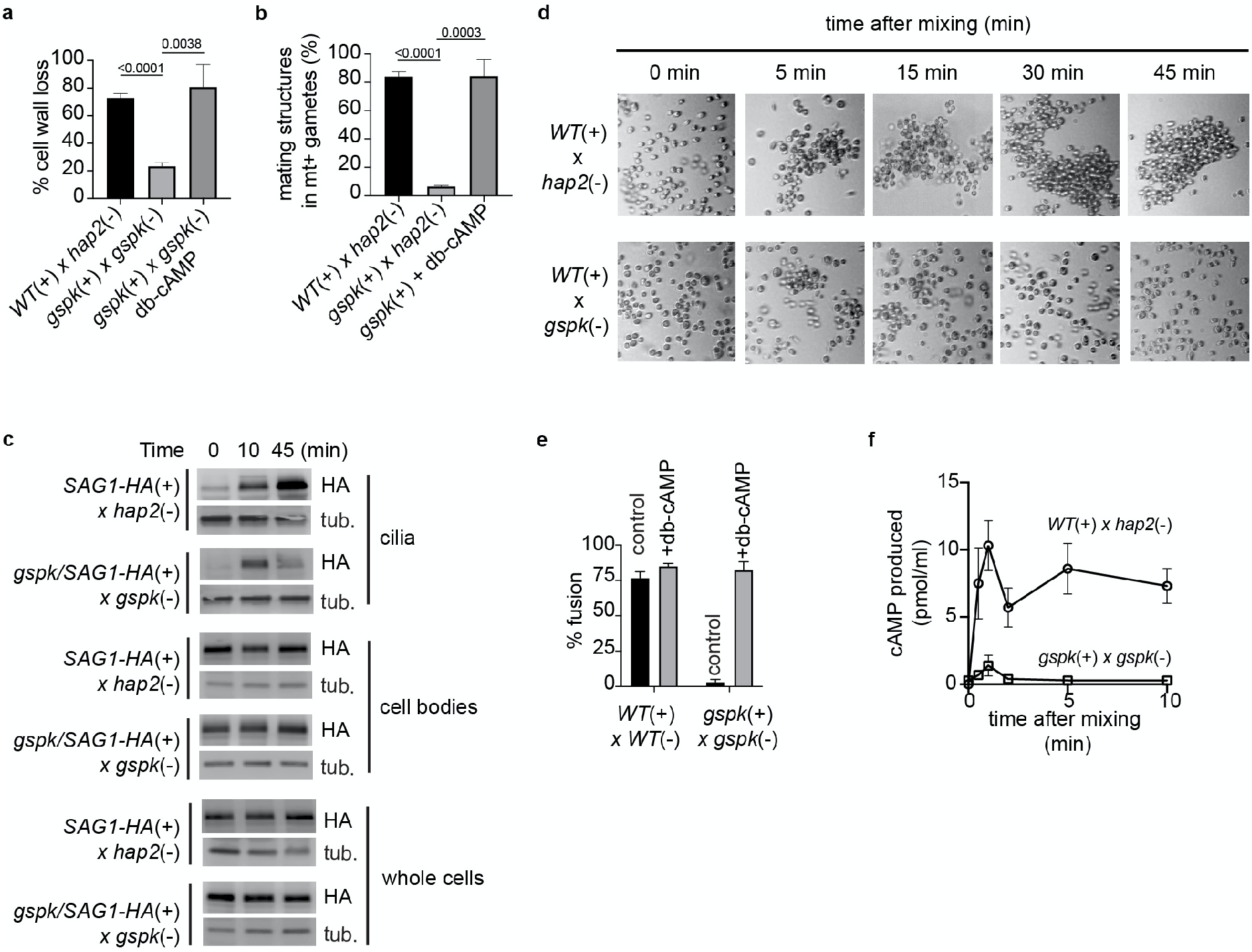
Ciliary adhesion-induced cell body responses are impaired in *gspk* mutants and restored by treatment with db-cAMP, and ciliary adhesion fails to trigger increases in cAMP in the mutants. **a-b** *gspk* gametes are impaired in cell wall loss and mating structure activation. Quantification of cell wall loss at 10 minutes after mixing *WT*(+) and *hap2*(-) gametes and *gspk*(+) and *gspk*(-) gametes in the absence and presence of db-cAMP buffer (**a**). Quantification of mating structure formation in mixtures of *WT*(+) and *hap2*(-) gametes and *gspk*(+) and *gspk*(-) gametes at 10 minutes after mixing and in samples of *gspk*(+) gametes that had been incubated with db-cAMP buffer for 30 minutes (**b**). **c** Ciliary adhesion-induced movement of SAG1-HA from the cell plasma membrane to the ciliary membrane is impaired in *gspk*(+) gametes. Anti-HA immunoblots of whole cells, cell bodies, and cilia from the indicated samples harvested 0, 10 and 45 minutes after SAG1-HA-expressing *WT* and *gspk*(+) gametes were separately mixed with *gspk*(-) gametes. The lower panel is a tubulin loading control. 3 μg protein were loaded in each lane. **d** *gspk* gametes fail to undergo sustained ciliary adhesion. Bright field micrographs of samples taken at the indicated times after mixing *WT*(+) gametes with *hap2*(-) gametes and *WT(+)* gametes with *gspk*(-) gametes. **e** Fusion in *gspk* gametes is rescued by db-cAMP. Quantification of fusion in mixtures of *WT plus* and *minus* gametes and *gspk plus* and *minus* gametes treated with or without db-cAMP. **f** The ciliary adhesion-induced increase in cAMP is impaired in *gspk* gametes. cAMP concentrations were determined by use of an ELISA-based method at the indicated times after mixing wild-type *plus* and *minus* gametes and *gspk plus* and *minus* gametes. P-values shown for A and B are from comparisons of means by Student’s t-test.

We also investigated the ability of *gspk plus* gametes to recruit SAG1 from the cell body to the cilia, a response to ciliary signaling that maintains and enhances ciliary adhesion^45^. We obtained *gspk plus* cells bearing HA-tagged SAG1 from a cross between *gspk*(-) gametes with *SAG1-HA*(+) gametes (Supplementary Fig. 2). *gspk*/SAG1-HA(+) gametes were mixed with fusion-defective *hap2*(-)gametes, and at 0, 10, and 45 minutes after mixing, samples were harvested, fractionated, and whole cells, cell bodies, and cilia were analyzed by anti-HA immunoblotting. Samples from a mixture of adhering *SAG1-HA* and *hap2* gametes served as controls. As expected, immunoblots of equal amounts of protein showed that SAG1-HA was present at low levels in cilia compared to the cell bodies in the 0-time samples of both the *WT* and *gspk*(+) gametes (Fig. 4c). Moreover, at 10 minutes after mixing, SAG1-HA had been recruited into both the *WT* and the *gspk* cilia. On other hand, at 45 minutes after mixing the amount of SAG1-HA had increased in the cilia of the *WT plus* gametes, but the amount of SAG1-HA in the cilia of the *gspk* gametes had decreased (Fig. 4c). Thus, although it was dispensable for the initial, adhesion-induced recruitment of SAG1-HA to cilia, GSPK was required to sustain SAG1 recruitment.

Ciliary adhesion is dynamic, and sites of adhesion are constantly being formed and broken along the lengths of the cilia as the organelles release membrane vesicles (ciliary ectosomes) enriched in SAG1 and SAD1^40^. Maintenance of ciliary adhesion, thus, depends on recruitment of SAG1 from the inactive pool on the surface of the cell body membrane. Indeed, consistent with the decreased amount of SAG1-HA in the cilia of the *gspk* gametes at 30 minutes after mixing (Fig. 4c), examination by phase-contrast microscopy of the 30 minute samples of the *gspk*/SAG1-HA *plus* gametes mixed with the fusion-defective *hap2 minus* gametes samples indicated that many had become single cells. And, by 45 minutes few if any were in clusters. On the other hand, in the control sample of *WT plus* gametes mixed with fusion-defective *hap2 minus* gametes, the cells continued to adhere to each other and the clusters had grown even larger (Fig. 4d). Thus, both biochemical and functional evidence indicated that sustained recruitment of adhesion molecules during ciliary adhesion required GSPK.

Taken together, the results above suggested that GSPK functioned upstream of all of the cell body events in the gamete activation pathway. Consistent with this interpretation, when db-cAMP was added to *gspk* gametes, they underwent cell wall loss and mating structure activation similarly to *WT* gametes (Fig. 4a, b). Furthermore, in the presence of db-cAMP, *gspk plus* and *minus* gametes were fully capable of undergoing cell-cell fusion (Fig. 4e), suggesting that GSPK was a positive regulator of cAMP. Indeed, assays for cellular cAMP showed that whereas the levels of this seond messenger increased nearly 10-fold when *WT plus* and *minus* gametes were mixed together, the cAMP increase was transient and less than 2-fold when plus and *minus gspk* mutants were mixed together (Fig. 4f). Thus, the rapid increase in cellular cAMP triggered by ciliary adhesion and required for gamete activation depends on this protein kinase located in the cytoplasm.

## Discussion

We screened for cell-cell fusion defects in several *Chlamydomonas minus* mating type strains from the CLiP mutant library annotated to have disruptions in gamete-specific protein kinase genes. Out of 9 strains, we identified one, with a mutation in the mixed lineage protein kinase gene *GSPK*, that underwent ciliary adhesion with *plus* gametes similarly to *WT minus* gametes but was strongly impaired in cell-cell fusion. *gspk* mutants were rescued for gamete fusion by introduction of a transgene encoding an HA-tagged form of GSPK, *GSPK-HA*. Cell fractionation and immunoblotting showed that GSPK-HA was present in cell bodies, with little if any in cilia. Immunoblotting in combination with treatment of cell lysates with a phosphatase enzyme showed that GSPK-HA was basally phosphorylated in naive gametes and that the entire cellular complement of GSPK was additionally phosphorylated within 1 minute after *plus* and *minus* gametes were mixed together. GSPK-HA was not phosphorylated when gametes were activated with db-cAMP, and db-cAMP treatment rescued fusion when added to *gspk plus* and *minus* gametes, indicating that phosphorylation of GSPK was not mediated by cAMP, and that GSPK functions upstream of the large increase in cAMP that induces gamete activation.

One of the most surprising findings was that all of the GSPK was phosphorylated within 1 minute after the gametes were mixed together (Fig. 2). Ciliary adhesion and consequent phosphorylation events indeed were terminated at the times indicated, because the samples for the immunoblots were placed directly into SDS-PAGE sample buffer and immediately heated. This response to engagement of receptors in cilia is much slower than that of olfaction, and slightly faster than that reported for the somatostatin receptor 3 pathway and the Hh pathway. In olfaction, the rapid increase in cAMP within the cilia of olfactory epithelial cells triggered by binding of odorants to their GPCRs leads to changes in plasma membrane potential detectable within 100 ms^46^, nearly 3 orders of magnitude more rapid than for the GSPK response.

In the somatostatin pathway, mobilization of cytoplasmic β-arrestin 2 into cilia was detected by immunofluorescence within 4 minutes after addition of somatostatin to cultured hippocampal neurons^47^, a response time of similar magnitude, but slower than the GSPK response. In the Hh pathway, increases in full-length forms of Gli transcription factors within cilia were detected by immunofluorescence within 5 minutes after addition of the Hh ligand to cells in culture^48^. In more recent reports, increases or decreases in ciliary levels of several other Hh pathway proteins, including soluble and transmembrane proteins, were detected ∼15 minutes after addition of the Hh ligand^49,50^.

One important difference between the GSPK and β-arrestin 2 responses and the Hh pathway response is that the first two depend on a signal sent from the cilium to the cell body, whereas the initial Hh pathway responses do not require communication between the cilium and the cell body but occur entirely within the cilium^16^. In the Hh pathway, Smo and the full-length Gli proteins are thought to move into and out of the cilium constitutively^48,51,52^. Through multiple, complex, and still emerging mechanisms, Hh binding to Patched activates Smoothened in the cilium, leading to Smoothened retention and consequent intraciliary alterations of Gli properties^12,52-55^. The Hh-dependent changes in cell proliferation that are the ultimate outcome of Hh pathway activation occur relatively much later^56^.

Our results indicate that ciliary adhesion indeed generates a signal that is sent to the cell body to elicit the large increase in cellular cAMP. That signal leads to phosphorylation of GSPK, and GSPK is required for the cAMP increase; but the failure of db-cAMP to induce GSPK phosphorylation indicates that the signal from the cilia is not cAMP, as been earlier suggested^24^, and thus the signal remains unknown. Similarly, the rapid movement of β-arrestin from the cytoplasm to the cilia upon activation of the somatostatin receptor 3 (SSTR3) is proposed to be a response to an undefined signal from the cilium^47^.

Another consideration also argues against the notion that cAMP from *Chlamydomonas* cilia triggers the responses in the cell body. In addition to their critical signaling role in sexual reproduction, the two cilia drive motility. Under the control of cues from light shining on the channelrhodopsin-containing eyespots in cells, the beating of the two cilia can be differentially controlled to allow the cells to swim toward or away from the light and find favorable environments for photosynthetic growth. Although photoreceptor currents that regulate calcium concentrations are the primary controller of motiity^57^, cAMP also plays a role^58^. Our results that GSPK responds to non-cAMP-mediated signals from the cilia provides a solution to the potential problem that changes in light intensity experienced by gametes would activate them for cell fusion in the absence of a partner.

Results from a *Chlamydomonas* mutant with a phenotype similar to the *gspk* mutant raise ideas about the nature of the undefined ciliary signal. Gametes of the *imp-3* mutant which have a lesion in the PP2A3 phosphatase^59^, also undergo normal ciliary adhesion, but adhesion fails to increase cAMP and fusion is strongly impaired. The substrates for PP2A3 are unknown, but earlier work on *imp-3* mutant gametes suggested that this phosphatase functioned in the cilia, not the cell body^60^. Consistent with this earlier observation, immunofluorescence studies showed that PP2A3 was enriched in the proximal part of the cilia, just distal to the transition zone and some was also localized in the cell body^59^. One scenario for adhesion-induced gamete activation would be that SAG1-SAD1 interactions somehow modify ciliary PP2A3, which then moves to the cytoplasm to carry out its function, which could include changing the phosphorylation state of proteins that regulate GSPK properties. Future experiments with gametes bearing combinations of *WT* and mutant forms of GSPK and PP2A3 should provide new insights into the nature of the signal transmitted from the cilia.

We should note that because of the rapid kinetics of the responses of *Chlamydomonas* gametes to ciliary adhesion, we cannot rule out the possibility that this signaling system is similar to those of cilium-based olfaction and vision and depends on changes in membrane potential. Indeed, *Chlamydomonas* possesses a gene, ADCY1/Cre06.g300500, that encodes an unusual chimeric protein with a predicted N-terminal channel-like domain and a C-terminal adenylyl cyclase domain. Moreover, our gamete transcriptome results^43^ showed that ADCY1 transcripts are gamete-specific and upregulated during gamete activation. ADCY1 homologs are present in ciliated protozoa where they regulate motility^61,62^.

A possible clue about mechanisms for movement of a signal from the cilium to the cell body comes from earlier work on the role of IFT in ciliary signaling in *Chlamydomonas*. Studies with gametes of the *fla10* temperature-sensitive mutant of the anterograde IFT motor kinesin-2 (FLA10) showed that *Chlamydomonas* gametes whose cilia were transiently depleted of their IFT machinery exhibited a ciliary signaling phenotype identical to the *gspk* phenotype^33,34^. At 45 minutes after *fla10* gametes were transferred to the non-permissive temperature, IFT components were depleted from the cilia, but the cilia remained essentially full-length and were undiminished in their ability to undergo ciliary adhesion. Importantly, though, ciliary adhesion failed to induce the typical increase in cAMP and failed to induce gamete activation and cell fusion. As with the *gspk* mutants, gamete activation was rescued by addition of db-cAMP. Thus, the signal for GSPK responses could be carried by retrograde IFT from the cilia to the cytoplasm. It will be interesting to determine whether IFT is also required for the SSTR3 and other GPCR responses in vertebrates. Unfortunately, the inability to conditionally deplete vertebrate cilia of their IFT machinery makes such experiments challenging. Conventional mutations in IFT proteins block ciliogenesis and conditional IFT mutants are only just becoming available^63^.

Notably, not only is the cilium-to-cytoplasm signal in the somatostatin pathway undefined, but (with the exceptions of odorant receptors, rhodopsin, and smoothened), the cellular and molecular mechanisms that link ligand binding by the multitude of other vertebrate ciliary GPCRs to responses in the cytoplasm remain largely unknown. Current models are that cAMP from the cilium diffuses into the cytoplasm to regulate effectors in the cytoplasm. But, whether cilium generated-cAMP that diffuses into the cytoplasm indeed is the signal is uncertain, and the localization and trafficking of effectors are still in early stages of investigation.

Our findings now set the stage for learning more about mechanisms of receptor-mediated cilium-to-cytoplasm communication. It will be important to learn whether, as we expect, the adhesion-induced phosphorylation of GSPK is essential for its function during gamete activation, and whether its protein kinase activity is required to induce the increase in cellular cAMP through as yet unidentified adenylyl cyclases or phosphodiesterases. Perhaps of even more importance, though, will be to use this system to investigate the undefined signal transmitted from the cilium to the cytoplasm and the mechanism of its transport.

## Acknowledgments

We are grateful to Dr. Caren Chang, University of Maryland, College Park, MD, USA for insightful discussions. We thank our laboratory colleagues, Drs. Jennifer Pinello and Jun Zhang for their constructive insights. We acknowledge the Imaging Core Facility in the department of Cell Biology and Molecular Genetics at the University of Maryland, College Park for Leica TCS SP5 confocal microscope. This work was supported by National Institutes of Health Grant GM122565 to W. J. S.

## Author Contributions

Conceptualization: M.A., P.R., W.J.S. Investigation: M.A., P.R., W.J.S. Methodology: M.A., P.R., P.H., S.K., W.J.S. Resources: M.A., P.R., P.H., S.K., W.J.S. Writing:, M.A., P.R., S.K., W.J.S. Reviewing: M.A., P.R., S.K., P.H., W.J.S.

## Methods

### Contact for Reagent and Resource Sharing

Requests for further information or resources and reagents should be directed to and will be fulfilled by the Lead Contact, William J. Snell.

### Cells and cell culture

*Chlamydomonas reinhardtii* wild type strains *21gr* (mating type *plus*; mt+; CC-1690; design*at*ed *WT*(+), *CMJ030* (mating type *minus*; mt−; CC-5325; designated *WT*(-), *hap2 (40D4;* CC5281) and *SAG1-HA* strains used in this study were grown in liquid tris-acetate phosphate medium (TAP) medium containing trace metals) at 22°C with aeration, or on the TAP plates with 1.5% agar^36^. The *Chlamydomonas* CLiP library mutants were obtained from the *Chlamydomonas* Resource Center. These mutants were generated by the insertion of a DNA cassette (CIB1) conferring resistance to paromomycin into the *Chlamydomonas* strain CMJ030^64^. Upon receipt, each mutant was streaked to single colonies, genomic DNA was isolated using Clontech plant genomic DNA isolation reagent (Takara, Cat. No. 9194), and the insertion site of the CIB1 cassette was verified by PCR. The PCR primers used to confirm the insertions are listed in Supplementary Table 1.

### Plasmid construction and transformation into *Chlamydomonas*

To prepare a plasmid containing an HA-tagged *GSPK gene*, a gene fragment of 8780/8769 bp that included the full-length *GSPK* gene sequence (7272 bp) and an additional 850 bp 5’ to the annotated transcription start site predicted to include the endogenous promoter and an additional 647 bp 3’ to the stop codon was amplified from DNA of BAC clone 34G21 by PCR using primers possessing *Xho1* and *Not1* restriction sites at the 5’ and 3’ ends, respectively. The amplified PCR product was cloned into a paromomycin resistance vector, *pChlamiRNA3int* (obtained from the *Chlamydomonas* Resource Center) in between *Xho1* and *Not1* restriction sites by In-fusion HD EcoDry cloning plus kit (Takara, Cat. No. 638915). A gene fragment encoding three copies of the 9-amino-acid HA epitope followed by *EcoR1* and *XbaI* restriction sites was inserted using QuikChange II XL Site-Directed Mutagenesis Kit (Agilent technologies). The resulting *GSPK-HA* transgene plasmid (13,468 bp) containing the paromomycin resistance gene was verified by sequencing. For *Chlamydomonas* transformation, *pGSPK-HA* was linearized with *BspH1* and the purified, linear plasmid was electroporated into *21gr* mt+ and CC-5325 mt-*Chlamydomonas* strains^65^. Transformants that grew on TAP plates containing paromomycin (Sigma, Cat. # P5057) were picked into 96-well plates and screened for the presence of *GSPK-HA* by PCR using primers listed in Supplementary Table 1. PCR-positive transformants were screened for GSPK-HA expression by immunoblotting with an anti-HA antibody.

### Gametogenesis, gamete activation, cell adhesion, and gamete fusion

Gametogenesis was induced by transferring vegetatively growing cells from TAP medium into N-free medium followed by growth under continuous light with aeration or agitation overnight. For gamete activation experiments, *plus* and *minus* gametes were mixed together or gametes of single mating types were experimentally activated by incubation in N-free medium containing 15 mM db-cAMP and 150 μM papaverine (db-cAMP buffer) for ∼10 minutes or more^22^. Cell-cell adhesion was quantified using an electronic particle counter (Coulter, Palo Alto, CA) as described previously^39,66^. Assays for cell wall loss and gamete fusion were as described previously^44,67^.

### Cell fractionation, cilia isolation, and assaying PKG phosphorylation

Fractionation of the cells into cell bodies and cilia from naive and adhering gametes was carried out as described earlier^36^. Phosphorylation of PKG was assayed in vitro as described earlier^36^ using a protein tyrosine kinase (PTK) assay. 20 μl of whole cilia (∼3 μg/μl protein) in 5% sucrose, 20 mM HEPES buffer were mixed with 20 μl of 2X PTK buffer (20 mM HEPES, pH 7.2, 10 mM MgCl2, 2 mM dithiothreitol, 1 mM EDTA, 50 mM KCl, 2 mM ATP, 0.2% Nonidet P-40, 0.4 mM orthovanadate, 20 mM β-glycerolphosphate, and 2% Sigma plant protease inhibitor cocktail) in the presence of ATP for 10 minutes and the phosphorylated form of PKG was detected by use of 4-20% gradient SDS-PAGE gels and immunoblotting using anti-phospho-tyrosine (anti-p-Tyr) antibody (Sigma, Cat. # 05-321).

### GSPK phosphorylation and λ-phosphatase treatment

GSPK phosphorylation in lysates of naive, adhering, or db-cAMP-activated gametes was asssed by changes in migration in immunoblots. The reactions were stopped by addition of aliquots to 4xSDS sample buffer followed by immediate boiling. For phosphatase treatment, the lysates was prepared by brief sonication of 2×10^7^ cells/ml in 1 ml in HEMDK buffer. The 40 μl final reaction volume contained 31 μl cell lysate, 1 μl λ-phosphatase (NEB, 400,000 U/μl), and 8 μl of phosphatase reaction buffer (NEB). After incubation at 30°C for 30 minutes, reactions were terminated by adding 40 μl of 4 x SDS sample buffer followed by boiling. As controls, samples were incubated in the presence of a phosphatase inhibitor cocktail (Sigma Cat. No. P2850).

### cAMP ELISA assay

cAMP levels in adhering wildtype and *GSPK* mutant gametes were quantified by use of a cAMP Elisa kit (Enzo Life Sciences, #ADI-900-163). Equal numbers (100 µl, 2×10^7^ cells/ml in M-N) of the *WT* and *gspk plus* and *minus* gametes were separately mixed in 1.5 ml Eppendorf tubes to initiate ciliary adhesion, and at the times indicated the cells were harvested by centrifugation (6350 x g; 4°C) and flash-frozen in liquid nitrogen. For the assay, samples were resuspended in 100 µl of 0.1 M HCl and incubated at room temperature for 10 minutes followed by clarification by centrifugation (20000 x g; 4°C). Supernatants were transferred to fresh tubes for use in the assay, which was performed using the acetylation protocol according to the manufacturer’s instructions. Absorbance at 405 nm of standards and experimental samples were determined using a microplate reader (LabSystems-Multiskan Ascent 354 Microplate Reader, San Diego, CA, USA). Results shown are from 6 independent experiments and are plotted as pmol/ml cAMP produced in the *WT* and *gspk* mutant mixtures.

### Determination of mating structure activation

As described previously^68^, samples (∼200 μl, 5×10^6^ cells/ ml) in N-free medium were seeded on cover slips coated with poly-L-lysine (Sigma, Cat. No. P8920) for 5 minutes followed by fixation with 4% paraformaldehyde solution (Sigma, Cat. No. 158127) freshly made in 10 mM HEPES, pH 7.4. Coverslips were washed with 1x PBS for 3 minutes, immersed for 6 minutes in 80% acetone (Fischer Scientific, Cat. No. 67-64-1), 30 mM NaCl, and 2 mM sodium phosphate buffer, pH 7.0, at −20°C, followed by immersion in 100% acetone at −20°C for 6 minutes. Samples were then stained with Allexa 488 Phalloidin (ThermoFisher Scientific, Cat. No. A12379) for 15 minutes in the dark^69^. Phalloidin incubation was followed by a wash in 1x PBS for 5 minutes. Finally, coverslips were mounted on slides using antifading agent Fluoromount-G™ (ThermoFisher Scientific, Cat. No. 00-4959-02) and examined by Hyd detector-equipped Leica TCS SP5 confocal microscope using a 1.4 numerical aperture, 63 X oil immersion objective. Images obtained from z series were summed to produce a projected image using Leica LAS X, and cropped in the Illustrator program of Adobe Systems (USA).

### Protein Determination, SDS PAGE and Immunoblotting

Protein concentrations were determined by use of the Bradford assay (Bio-Rad protein assay kit II, Cat. No. 5000002). For immunoblotting, samples were separated by SDS-PAGE on 4–20% Tris-Glycine or SDS-MOPS gradient gels (GenScript, USA) and transferred onto PVDF membranes (Merck Millipore, Cat. No. IPVH00010) as described previously^34,40^. Membranes were blocked by incubation in 3% fat-free dried milk (Carnation, Nestle, Inc., Solon, OH) for 1 hour followed by 1 hour of incubation in the primary antibody. Membranes were washed three times for 10 minutes with TBST (Tris-buffered saline, 0.1% Tween 20) followed by incubation with secondary antibody. After three consecutive washes with TBST and incubation in the chemiluminescent substrate, fluorescence signals were captured on a C-Digit blot scanner (LI-COR Instruments, USA). The antibodies used for immunoblotting were rat anti-HA (1:3000; Roche), mouse anti-α-tubulin (1:5000; Sigma) and goat anti-rat IgG HRP(1:5000; Merck); and goat anti-mouse IgG HRP (1:5000; Sigma).

### Bioinformatic Analysis, Quantification and Statistical Analysis

For comparative analysis the homologous protein sequences were aligned with ClustalW, and the percentage of positions with identical or identical plus similar amino acid positions were calculated using the BioEdit 7.2 software (https://bioedit.software.informer.com/7.2/) software with a threshold of 75%. JPred4 was used to predict secondary structure^70^. N-terminal myristoylation (N-myristoylation) sites were predicted using NMT - The MYR Predictor (http://mendel.imp.ac.at/myristate/SUPLpredictor.htm). All quantitative data represent at least three independent sets of experiments. Statistical significance of differences between groups was assessed by Student’s t-test. Data were analyzed using GraphPad Prism 9 (GraphPad Software, U.S.A.).

## Supplementary information

**Supplementary Table 1:**
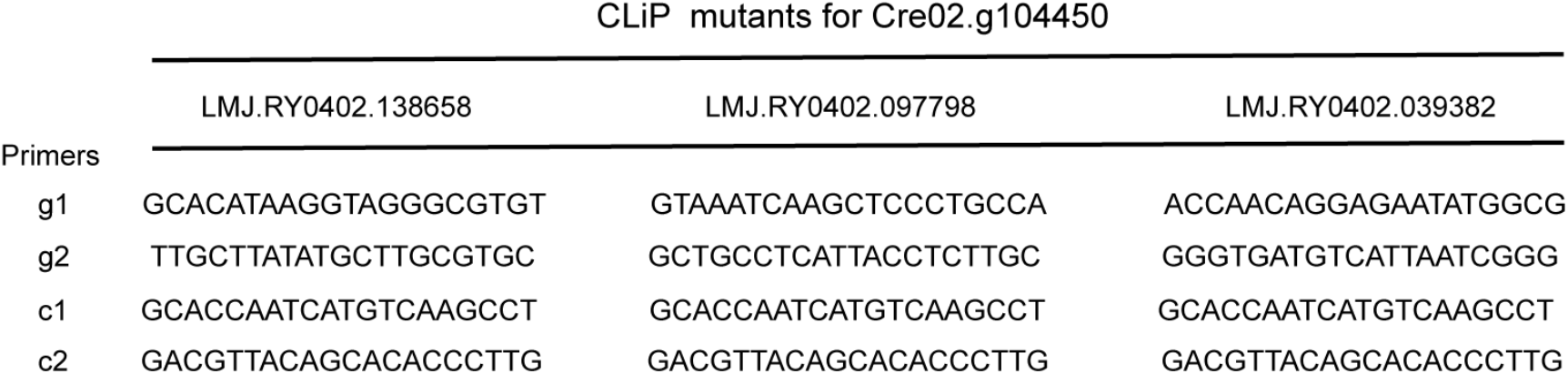
*AphVII* cassette and gene-specific primers used for CLiP mutant analysis.

**Supplementary Fig. 1:**
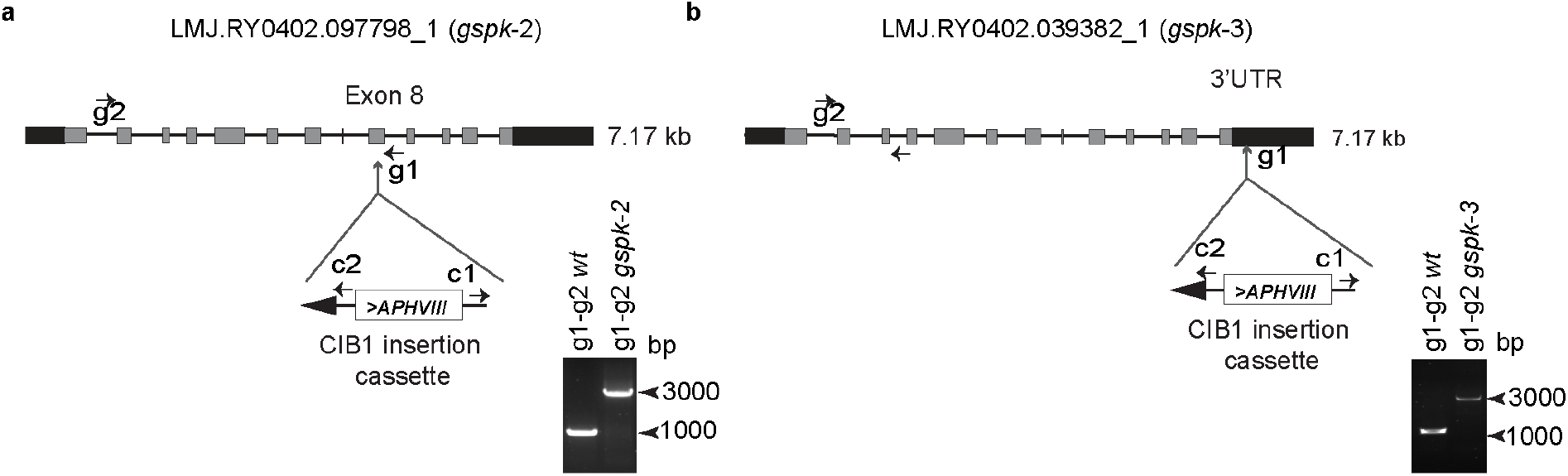
Structure of the *GSPK* genomic locus in CLiP library mutants. **a-b**. Grey solid boxes indicate exons; thin lines indicate introns; UTRs are shown as black boxes. The ClB1 (*AphVIII)* insertion cassette is located in the 8th intron of *gspk* in the *gspk* allele of LMJ.RY0402.097798_1 strain; *gspk-2* (**a**) and in the 3’UTR of *gspk* in the *gspk* allele of LMJ.RY0402.039382_1 strain; *gspk-3* (**b**). Primer locations for genotyping are shown by arrows where g1 and g2 are gene-specific primers and c1 and c2 are the primers specific to the insertion cassette. Gel images show the diagnostic genomic DNA PCR gels. The primer combinations used for PCR are indicated above the lanes. The PCR product of ∼1000 bp in the *WT Chlamydomonas* strain and the PCR products of ∼3000 bp in the *gspk-2* and *gspk-3* mutant strains document the *AphVIII* insertions.

**Supplementary Fig. 2:**
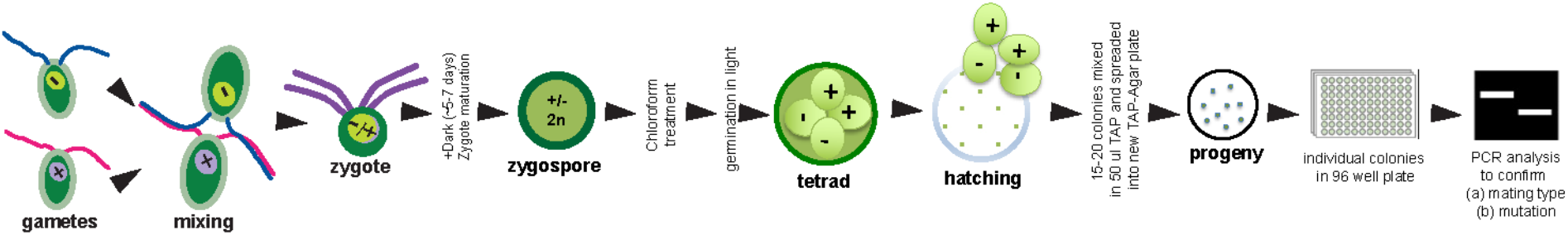
Graphical overview of method for obtaining progeny of desired genotypes from crosses of *Chlamydomonas* gametes.

**Supplementary Fig. 3:**
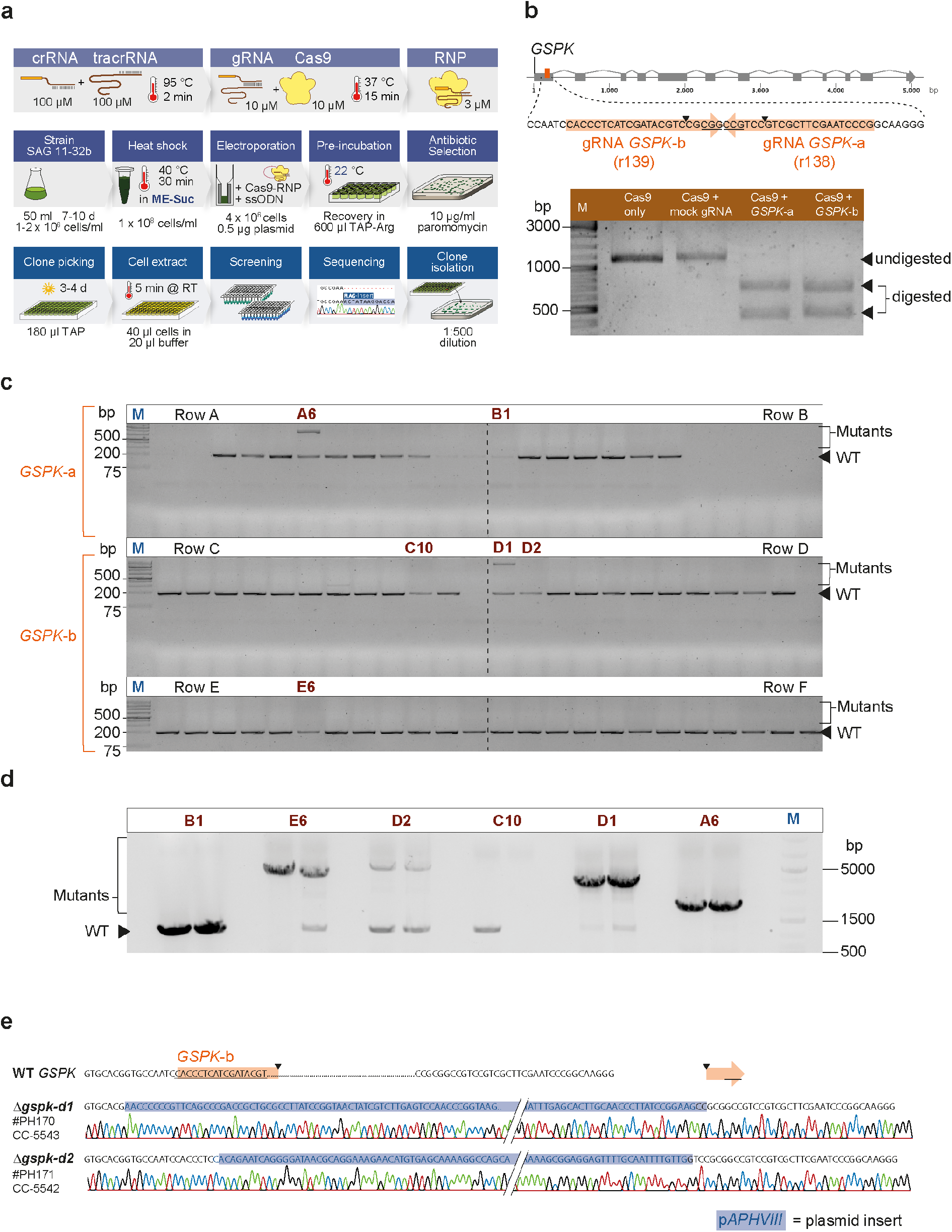
Generation of Δ*gspk-d2* CRISPR mutant. **a**. Graphical overview of the methodological steps for the generation of a *GSPK* mutant with CRISPR-Cas9 ribonucleoproteins (Cas9-RNPs). Taken and adapted from (Kelterborn, 2020) **b**. *GSPK* gene locus with two Cas9 target sites selected with the CRISPOR algorithm (Concordet and Haeussler, 2018). Efficiency of the two selected guide RNAs (gRNAs) was tested by *in-vitro* digestion as described in (Kelterborn, 2020). A PCR product spanning both target sites was incubated with only Cas9 protein, with Cas9-RNPs using a non-targeting mock gRNA, or with Cas9-RNPs assembled with *GSPK*-a or *GSPK*-b gRNA. The two lower DNA bands indicate effective Cas9-induced cleavage at the *GSPK*-a and *GSPK*-b target site. **c**. Single colonies transformed with *GSPK*-a or *GSPK*-b Cas9-RNPs were analyzed with a short colony-PCR spanning 197 bp containing both target sites. PCR bands with a different size (clone A6, C7 and D1) or different intensity than *WT* bands (clones A1, A2, A11, A12, C10, C12, D2, D10, E6) potentially indicate a mutated *GSPK* locus. **d**. Clone A6, B1, C10, D1, D2, E6 were selected for further analysis with a larger locus PCR (1157 bp) and using longer elongation times (3 min). PCR analysis reveal PCR bands with *WT* size (B1), large insertions (A6, D1, D2 and E6) or missing PCR bands (C10). A lower PCR band (∼1100 bp) can be seen in E6, D2, C10 and D1, and potentially derives from a mixture of mutant and WT cells. **e**. Clone Δ*gspk-d1* and Δ*gspk-d2* were singled out to remove remaining WT cells and the mutation in the *GSPK* locus was confirmed by sequencing analysis. Both clones show large insertions of the p*APHVIII* marker plasmid leading to a premature stop codon and consequently to disrupted *GSPK* gene expression. Cell-cell fusion results are shown for clone Δ*gspk-d2* CRISPR mutant.

**Supplementary Fig. 4:**
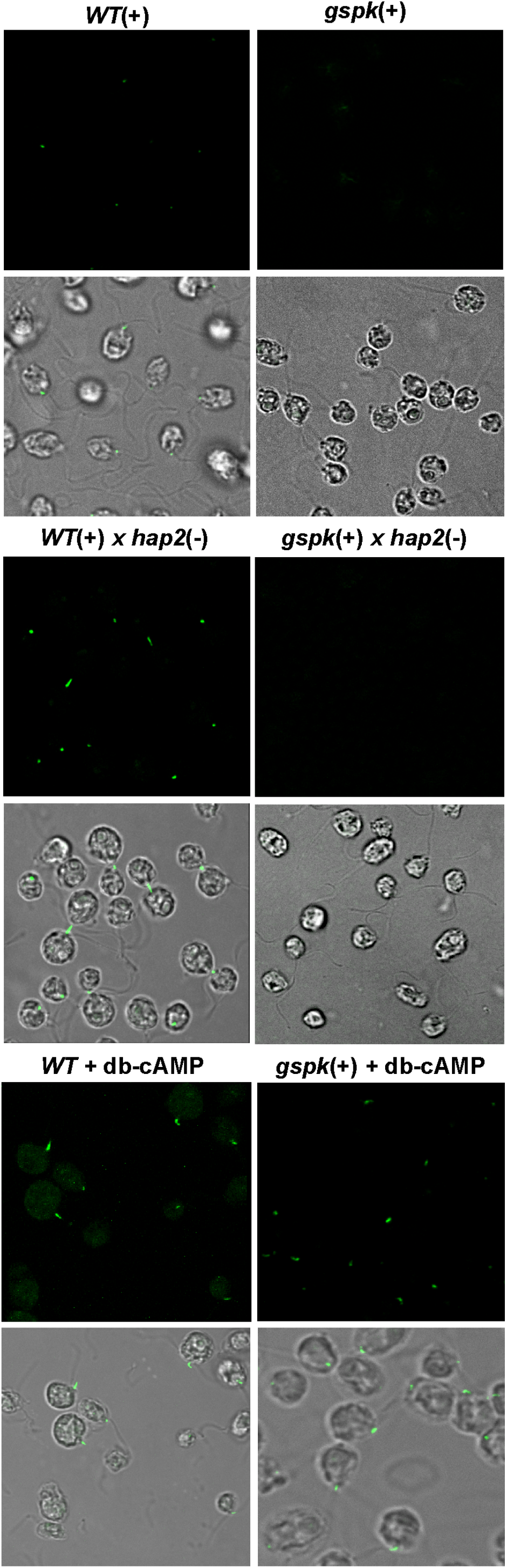
Ciliary adhesion induces formation of actin-filled fertilization tubules in *WT*(+) gametes but not in *gspk plus* gametes. Fluorescence images of cells stained with Alexa 488 phalloidin showed fertilization tubules in mixtures of *WT*(+) and *hap2*(-) gametes, but not in mixtures of *gspk*(+) and *hap2*(-) gametes (upper panel). Unmixed *WT*(+) and *gspk*(+) gametes lack fertilization tubules (control, middle panel). *gspk*(+) gametes activated with db-cAMP buffer formed fertilization tubules similarly to *WT*(+) gametes (lower panel).

